# Effects of planting density on the growth and photosynthetic characteristics of *Alternanthera philoxeroides* under different nutrient conditions

**DOI:** 10.1101/349860

**Authors:** Ke Li, Hao Chen, Haijie Zhang, Miansong Huang, Quan Quan, Tong Wang, Jian Liu

## Abstract

Density and nutrient level are important factors that might affect the growth of invasive plants. To reveal the effects of plant density on the performance of invasive plant *Alternanthera philoxeroides* under different nutrient conditions, a greenhouse experiment was conducted in which *A. philoxeroides* was planted at three densities (low, medium and high) under three nutrient levels (low, medium and high). The results showed that both planting density and nutrient levels had significant effects on the growth of the plant. The biomass of individual plant and all plants in one pot under medium nutrient level were the highest while the photosynthetic rate and total chlorophyll content were the highest at the high nutrient level. Under different nutrient levels, the photosynthetic rate was the highest at medium planting density. The biomass of single plant decreased with the increase of population density, while the total biomass in the whole pot increased with the increase of density. These characteristics might contribute to the invasion of *A. philoxeroides* and help the plant to form monodominant community.

## Introduction

Biological invasion has far reaching impacts on ecological functions and the ecological diversity of native environments [1-5]. The invasive species have high fecundity, and once the invasion succeeds, it is easy to form the predominant population. The success of biological invasion is not only determined by the physiological and genetic characteristics of invasive species, but also by the factors such as population density and nutrient levels [6-10].

The nutrient is one of the important factors for plant growth. Previous studies have found that traits associated with the resource used determine the invasion success of several invasive species [11, 12]. Invasive plants generally show stronger photosynthesis and growth when invading habitats with richer nutrients [12-15]. Therefore, the nutrient level is also associated with the invasion success [9]. The research on the influence of nutrient on invasive plants plays an important role in understanding the successful plant invasion.

The competition among plants is also an important factor affecting the spatial distribution, dynamics and species diversity, promoting the succession of communities [16]. Invasive plants can replace local plants through interspecific competition, accomplishing the process of successful invasion [17]. However, when invasive plants spread to a certain extent, there is a degree of intraspecific competition. In order to adapt to this change, plants tend to change their own characteristics, such as plant height, biomass of branches and physiological characteristics of leaves, etc [18, 19]. And this phenotypic plasticity allows invaders to allocate more nutrients than their native counterparts to increase biomass [20, 21]. Several studies found that the physiological indexes of invasive plant under high density are less than those under low population density [8, 22]. Plant density determines the competitiveness of aquatic clonal plants in complex habitats [23]. Therefore, we can analyze the growth mechanism by studying the morphological changes of plant. At the same time, the effect of population density on invasive plants is one of the core problems in the study of invasion ecology at present [24].

Aquatic species might have a fast response to nutrient enrichment, increasing their biomass rapidly, which is particularly true in the aquatic invasive species [9, 25-27]. Therefore, aquatic invasive species affects the productivity and management of land and water resources worldwide [28]. *A. philoxeroides* is one of these aquatic invasive plants. It is a clone weed that is native to South America and it is a stoloniferous and rhizomatous perennial herbaceous plant [28-30], which propagates clonally and expands rapidly in both aquatic and terrestrial habitats [31-34]. *A. philoxeroides* has experienced an invasion history of more than 80 years in China and become one of the most harmful invasive species [34]. At present, there are many researches focused on the physiological characteristics of *A. philoxeroides*, its response to natural environmental factors, and biological and chemical control [10, 35-37]. Many studies have shown that if the plant has a high photosynthetic rate, and it usually grows and propagate rapidly [38, 39]. Moreover, the invasive capacity of invasive plants is closely related to the ability of photosynthesis, they often enhance its invasive ability by increasing the photosynthesis of ramets [40, 41]. Thus, studying the photosynthetic capacity of *A. philoxeroides* is essential for understanding the potential of invasive species and to develop appropriate control strategies.

Our previous studies have compared the growth of *A. philoxeroides* with native plants [10]. So in the present study, we conducted a greenhouse experiment which combined nutrient levels with planting density to explore their effects on the growth and photosynthetic characteristic of *A. philoxeroides*. We asked the following questions: (1) Will the increase of planting density inhibit the growth of *A. philoxeroides* under various nutrient levels? (2) How do the nutrient level and planting density affect the invasion of *A. philoxeroides*.

## Materials and methods

### Ethics statement

We collected plant material for our study with the official permission of the Environmental Protection Bureau of Weishan County, and the Management Committee of the Weishan Lake Constructed Wetland Park. We did not collect endangered or protected species.

### Plant material and experimental design

The *A. philoxeroides* seedlings were collected from Weishan Lake Wetland Park, Shandong province, in July 2017. The seedlings were cultured a week in the greenhouse of Fanggan Research Station of Shandong University (36°26’N, 117°27’E). The method of sand culture was used in the experiment, and the river sand was washed thoroughly before planting. The length of the selected stem was about 11cm, and then was transplanted into a pot (h: 23.5cm; d: 22cm) with 7 kg sand. Set up two factors in our experiment: the nutrient level of the sand and planting density of *A. philoxeroides*. The nutrient treatments consisted of three levels (low, medium and high; labeled as A, B and C) and three kinds of planting density (low, medium and high; 1, 2 and 4 seedlings per pot, respectively). In total, there were nine treatment combinations, each dealing with five replicates. The nutrient levels and planting density settings in the experiment are shown in Table 1.

**Table 1.** Nutrient levels and planting density of *A. philoxeroides* at different treatment.

| Nutrient levels | TN<br>(g kg<sup>-1</sup> sand) | Amount of chemical<br>fertilizer added<br>( g kg<sup>-1</sup> sand ) | planting density | planting number<br>(plant pot<sup>-1</sup>) |
| --- | --- | --- | --- | --- |
| A | 0.5 | 2.5 | 1 | 1 |
|  |  |  | 2 | 2 |
|  |  |  | 4 | 4 |
| B | 0.7 | 3.5 | 1 | 1 |
|  |  |  | 2 | 2 |
|  |  |  | 4 | 4 |
| C | 2.1 | 10.5 | 1 | 1 |
|  |  |  | 2 | 2 |
|  |  |  | 4 | 4 |

\*A, B, and C: low, medium and high nutrient levels.

In the experiment, the compound fertilizer with the ratio of nitrogen and phosphorus (N: P = 4: 1) similar to Nansi Lake was used as the nutrient source. According to our previous experiments, the medium nutrient gradient in this experiment is the most suitable for the growth of *A. philoxeroides* [37]. Water 200 mL every day, in order to ensure the normal growth of *A. philoxeroides*, and would not lead to the loss of fertilizers in the pot. At the time of fertilization, grounding the fertilizer into powder, then half dissolved in 200 mL of water each time, added for two days, to prevent once adding cause damage to plants. At the time of experiment, the low nutrient level was added the fertilizer only in the first week; medium and high nutrient levels were added every two weeks during the experiment. The time of the experiment was from the July 24, 2017 to September 20, 2017.

### Determination of photosynthetic rate and light response curve

The photosynthetic characteristics of each group were measured by a Portable Photosynthesis System (*LI-COR 6800*, USA) using PAR of 1000 µmolm^-2^s^-1^. Leaves were measured under ambient CO_2_ concentration [385 µmolmol^-1^] [42]. The light response curves were measured under the PAR of 1600, 1200, 1000, 800, 600, 400, 200, 100, and 0 µmolm^-2^s^-1^[42]. The fitting of the light response curve adopts a non - right - angle hyperbolic model, and the model formula is:

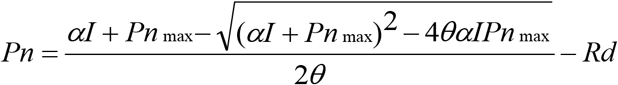

In this formula, α is the apparent quantum rate; *I* is the light quantum flux density; Pn_max_ is the maximum photosynthetic rate; Rd is the dark respiration rate; θ is the angle parameter that reflects the degree of bending of the light response curve, and the range of values is 0 ≦ θ ≦ 1 [43]. According to the light response curve, we calculated the light compensation point (LCP) and the light saturation point (LSP).

### Determination of chlorophyll content

In each treatment, using a volume fraction of 95% ethanol extract chlorophyll, then the spectrophotometer was used to measure the absorbance values at 649nm and 665nm wavelength [44]. The concentration of chlorophyll a, chlorophyll b and total chlorophyll were calculated according to the formula:

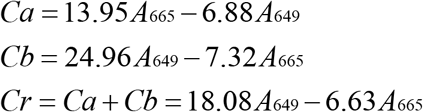

In the formula, Ca, Cb and Cr represent the concentrations of chlorophyll a, chlorophyll b and total chlorophyll respectively, [mg cm^-2^]; A_649_ and A_665_ represent the absorbance at the wavelengths of 649 nm and 665 nm respectively [45].

### Determination of specific leaf area (SLA) of *A. philoxeroides*

Before harvest, randomly selected 20 mature leaves of each treatment, wiped clean and flatted on the scanner. The determination of leaf area using Photoshop software.

Leaf area = the percentage of leaf pixels / background pixels · background paper area [46-48].

SLA = leaf area / leaf dry weight.

### Determination of morphological indexes of *A. philoxeroides*

We measured the length of stolon and recorded the number of internode before harvest.

The internode length of *A. philoxeroides* under different treatments was calculated.

### Determination of biomass index of *A. philoxeroides*

At the end of the experiment, the roots of *A. philoxeroides* were washed thoroughly, and the leaves, stems and roots of each treatment were respectively put into the envelopes, numbered and then dried up to constant weight in an oven which the temperature was 80 °C, recorded the dry weight of each part.

### Statistical analysis

The date of different variables, such as biomass, photosynthetic rate, SLA and total chlorophyll content was analyzed by two-way ANOVA with SPSS 21.0 software (Table 2). The significance test in all tests was performed at a level of P <0.05. Use Origin 8.5 software to draw charts.

## Results

**Table 2.**
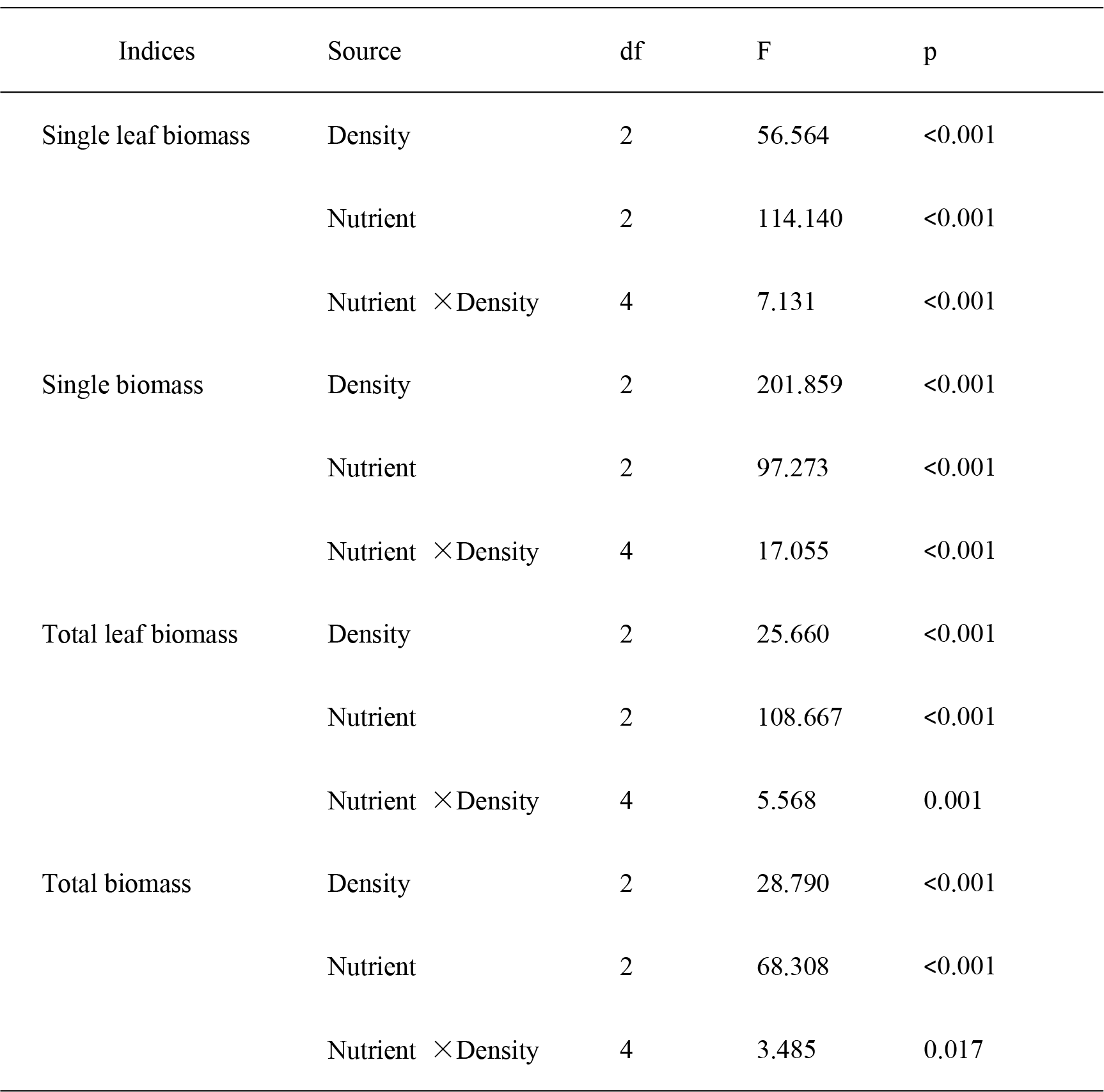
The two-way ANOVA of the effects of the independent variable nutrient level and plant density, and their combination on studied parameters of species *A. philoxeroides* in the experiment.

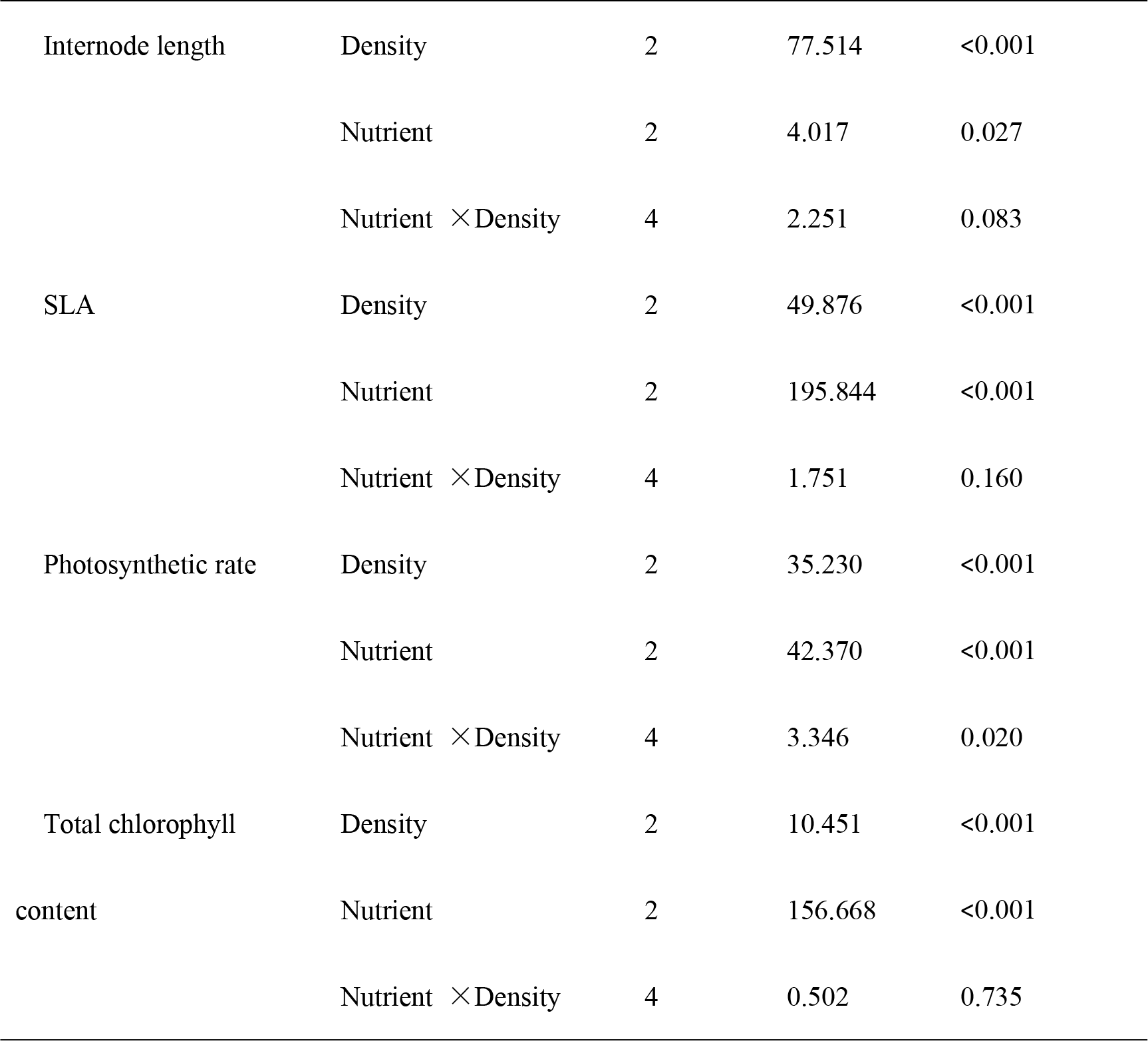

### Effects of density on Plant Biomass Index and Morphology Index under different nutrient gradients

Results showed that the interaction between nutrient level and plant density significantly affect the leaf biomass of per plant, single plant biomass, total leaf biomass and total biomass of a whole pot (Table 2).

Under three nutrient levels, the leaf biomass of single plant and the biomass of single plant all decreased significantly with the increase of planting density (Table 3). Among various nutrient levels, under three planting density, the reduction rate of leaf dry weight per plant was 47.9%, 53.9%, 62.5%; the reduction rate of biomass per plant was 57.2%, 64.4%, 63.4% (Table 3). At different nutrient levels, the difference in the leaf biomass of per plant between low density (1) and medium density (2) was slightly greater than the difference between medium density (2) and high density (4) (Table 3). At every nutrient levels, there was a significant difference in the biomass of single plant among all three planting densities (Table 3). In low nutrient level (A), the difference between medium planting density (2) and high planting density (4) is slightly larger than that between low planting density (1) and medium planting density (2) (Table 3). Under medium nutrient (B) and high nutrient (C), the gap between low density (1) and medium planting density (2) is slightly larger than medium planting density (2) and high planting density (4) (Table 3).

At the three nutrient levels, the total leaf biomass and the total biomass of a pot all increased significantly with the increase of planting density (Table 3). To total leaf biomass of *A. philoxeroides*, there was no significant difference between low planting density (1) and medium planting density (2) under the treatment of low (A) and medium (B)nutrient levels and the difference among the three planting densities was not significant under the high nutrient level (C) treatment (Table 3). For total biomass of a pot, at low nutrient level (A), the difference between medium planting density (2) and high planting density (3) is significantly smaller than that between low planting density (1) and medium planting density (2), besides there is no significant difference between low planting density (1) and medium planting density (2) at medium (B) and high (C) nutrient levels (Table 3).

According to the two-way ANOVA analysis (Table 2), there were have obvious interaction between nutrient level and plant density on internode length of *A. philoxeroides*.

With the increase of planting density, the average internode length of *A. philoxeroides* decreased in varying degrees, and the internode length was the longest at the treatment of low planting density (1) (Table 3). At the treatment of three nutrient levels, the average internode length of medium nutrient level (B) was the highest, is 5.6 cm (Table 3).

**Table 3.**
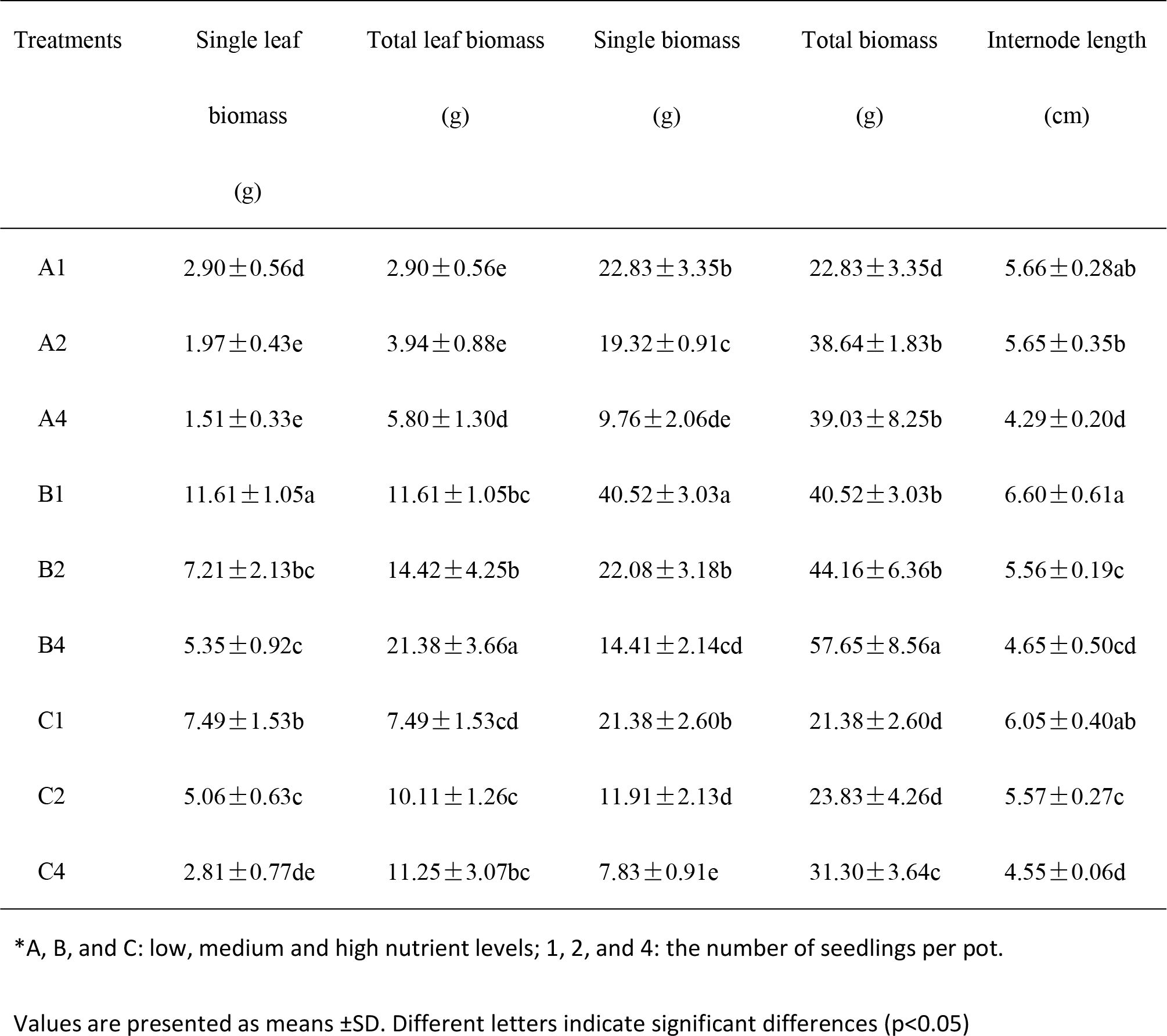
Different nutrient levels and different planting densities of *A. philoxeroides* biomass and morphological index.

### Effects of plant density on SLA of *A. philoxeroides* under different nutrient levels

According to the two-way ANOVA analysis (Table 2), there is no interaction between effects of nutrient level and plant density on SLA of *A. philoxeroides.*

The SLA of *A. philoxeroides* increased significantly with the increase of nutrient level (Fig 1). Different planting density had certain effects on the SLA of *A. philoxeroides*, which was the highest under medium planting density (2) and the lowest under high planting density (4), and both have significant differences (Fig 1).

**Fig 1.**
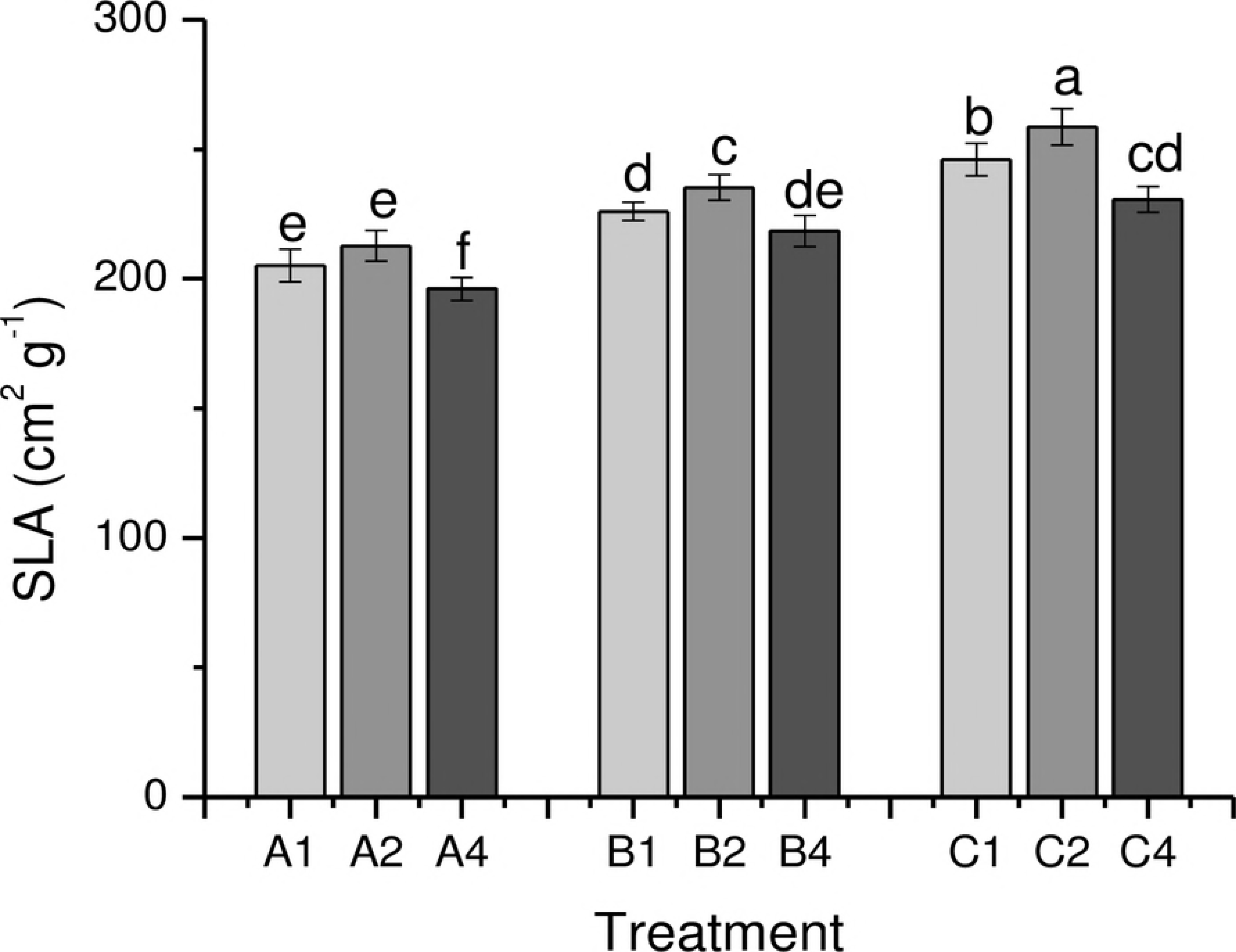
The SLA of *A. philoxeroides* at different nutrient level and plant density. A, B, and C: low, medium and high nutrient levels; 1, 2, and 4: the seedlings per pot. Values are presented as means ±SD. Different letters indicate significant differences (p<0.05).

### Effects of density on photosynthetic rate and total chlorophyll content of *A. philoxeroides* under different nutrient levels

We found that nutrient levels and planting densities on the photosynthetic rate of *A. philoxeroides* has a obvious interaction, and for the total chlorophyll content, they had no significant interaction to it (Table 2).

The analysis of photosynthetic rate and total chlorophyll content of *A. philoxeroides* under different treatments showed that the change trend of the two indexes are basically the same, and both of them increased with the increasing nutrient level (Fig 2). The photosynthetic rate of medium planting density (2) of the three nutrient levels were the highest and had significant differences, and they were the lowest under the treatment of high density (3) (Fig 2-A). The total chlorophyll content of the medium nutrient (B) and high nutrient (C) had obvious differences (Fig 2-B). The total chlorophyll content of *A. philoxeroides* reached the maximum under high nutrient level (C), and for the planting density, when the planting density was medium (2), the total chlorophyll content of *A. philoxeroides* reached the maximum (Fig 2-B).

**Fig 2.**
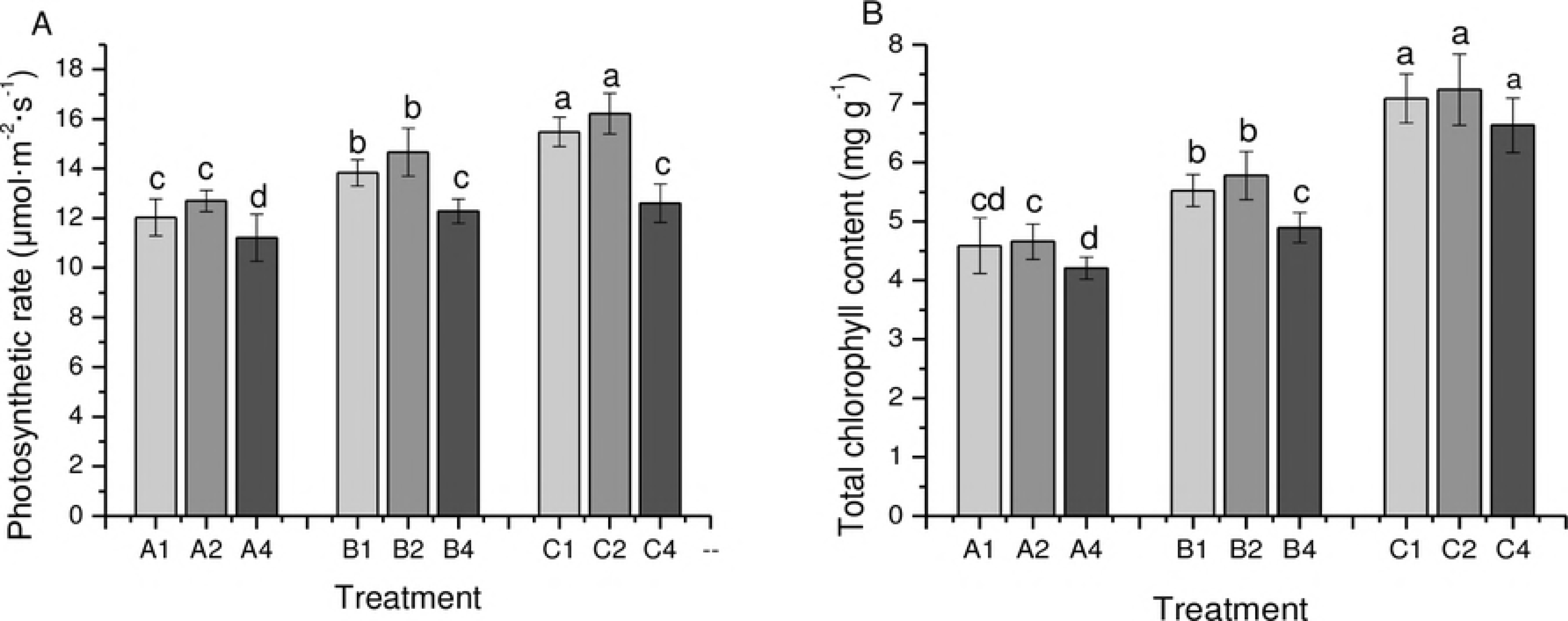
The photosynthetic rate and chlorophyll content of *A. philoxeroides* at different nutrient levels and plant densities. A, B, and C: low, medium and high nutrient levels; 1, 2, and 4: the number of seedlings per pot. Values are presented as means ±SD. Different letters indicate significant differences (p<0.05).

### Effects of density on light response curve of *A. Philoxeroides* under different nutrient levels

The data were analyzed and fitted by SPSS software. The fitting coefficients (R^2^) of the light response curve of each group were all greater than 0.9, which showed that the curve fitting degree was better and the photosynthetic characteristic of *A. philoxeroides* could be more accurately reflected.

With the increase of the nutrient level, the maximum photosynthetic rate (Pn_max_) of *A. philoxeroides* increased in different degrees, so the Pn_max_ at the high nutrient level (C) were the largest (Table 4). Among different planting density the Pn_max_ of *A. philoxeroides* were the largest at the treatments of medium planting density (2), and the Pn_max_ of *A. philoxeroides* was minimal under the high planting density (4) (Table 4). Besides, under the same nutrient level treatment, there were significant differences among the three different planting densities on the Pn_max_ of *A. philoxeroides* (Table 4).

The light compensation point (LCP) and light saturation point (LSP) of *A. philoxeroides* under different nutrient treatments were the biggest when planting density was medium (2), and when the planting density was high level (4), the value of LCP and LSP were the smallest (Table 4). At the same planting density, both LCP and LSP increased with increasing nutrient levels, among them, there was a larger gap between medium nutrient (B) and high nutrient (C) (Table 4). Under the same nutrient level, at the difference of the value of LCP and LSP between medium planting density (2) and high planting density (4) was larger than that between medium planting density (2) and low planting density (1) (Table 4).

**Table 4.**
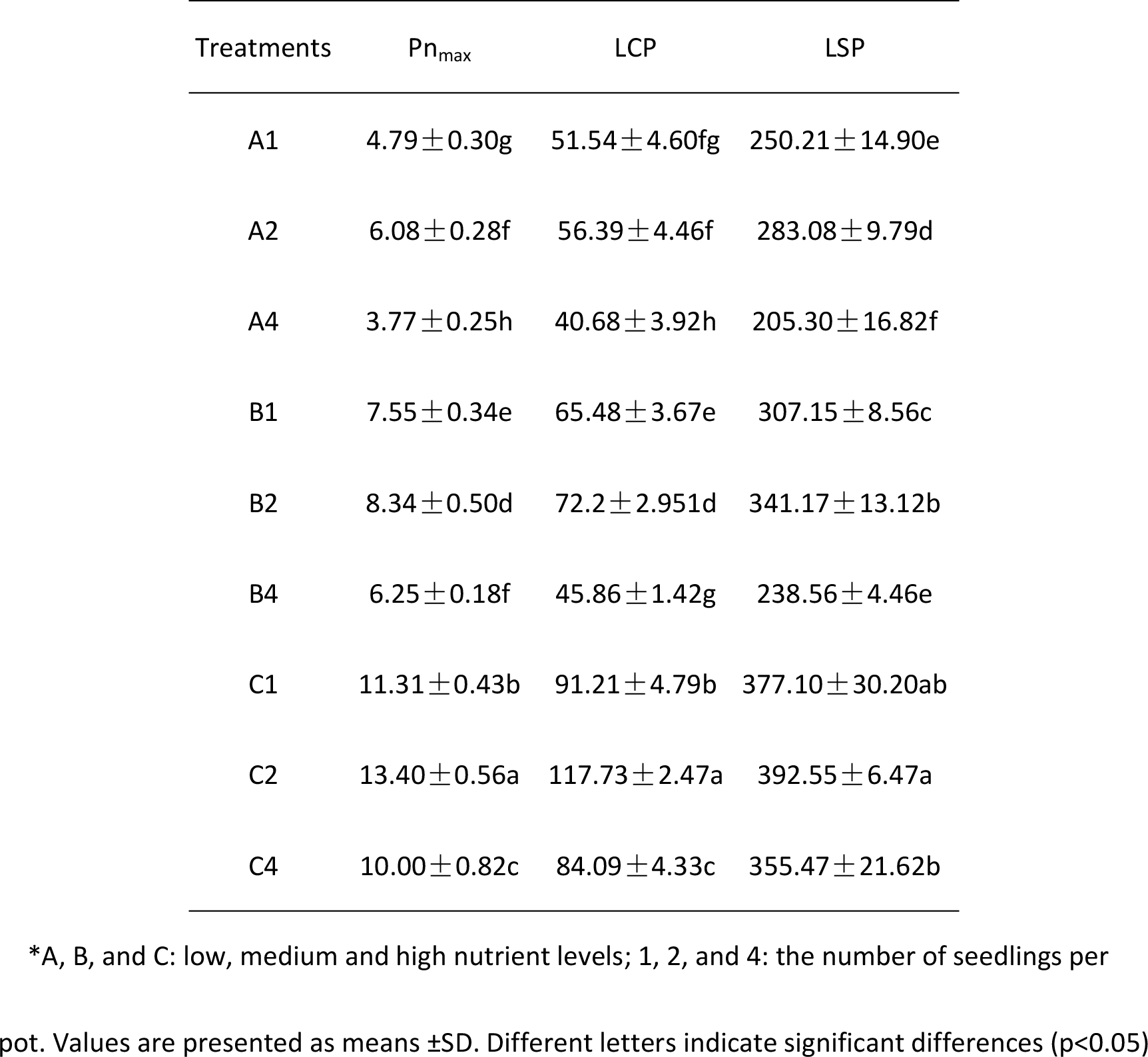
Light response curve parameters of *A. philoxeroides* at different nutrient levels and planting densities.

## Discussion

In our experiment, nutrient and planting density had significant interaction effects on the biomass accumulation of *A. philoxeroides*. Under the same planting density, the biomass accumulation of *A. philoxeroides* could be promoted by increasing nutrients [49-51]. And our previous studies concluded that compared with native plants, the *A. philoxeroides* had better environmental adaptability and higher biomass and ratio of leaf area under different nutrient conditions [10, 37]. At the treatment of medium nutrient level, the whole pot biomass of *A. philoxeroides* among three different planting densities was the highest, indicating that too high nutrient levels could be harmful for *A. philoxeroides* at the initial stage. At medium nutrient level, the increase of planting density had the most obvious effects on the single plant biomass of *A. philoxeroides* and the effects will be weakened at low nutrient. Therefore, we conclude that under appropriate nutrient level, the effects of planting density on the growth of *A. philoxeroides* were nutrient dependent. Some studies have shown that at same nutrient condition, the biomass of invasive plants increased more than that of native plants [52]. So in our experiments, under the same nutrient level, although the biomass of single plant was reduced with the increase of planting density, the total biomass of the plant in whole pot was increased, especially at low nutrient. That would enhance the invasive ability of *A. philoxeroides*.

In addition, according to the leaf biomass of *A. philoxeroides*, we analyzed the SLA of *A. philoxeroides*. In our study, with the increase of nutrient level, the SLA of *A. philoxeroides* increased, which is consistent with the studies on SLA of other plants [53-55]. Similarly, the increase of planting density has different influences on the SLA of *A. philoxeroides*. In this study, the SLA of *A. philoxeroides* was the highest under medium planting density treatment. SLA is one of the important plant leaf traits and closely related to plant growth and survival strategy. Its value can reflect the ability of plant leaves to intercept light and self-protection in bright light [56]. It is closely related to the photosynthesis and respiration of plants [57]. We therefore concluded that at the medium planting density, the leaf of *A. philoxeroides* had a higher net photosynthetic rate. So we then analyzed the photosynthetic rate of rate of each treatments to confirm our conclusion.

In this study, photosynthetic rate had increase with the increasing nutrient levels that is consistent with previous studies [58-60]. Among the three kinds of plant density treatment, the photosynthetic rate in medium planting density was the highest, and the difference with high planting density was obvious. Chlorophyll is the main pigment of photosynthesis in plants, which reflects the size of photosynthesis in plants [44]. By this experiment, we found that the content of the total chlorophyll of *A. philoxeroides* had the same trend, indicating that under medium planting density, plants can capture resources better [61].

In order to better understand the effects of planting density on the photosynthetic characteristics of *A. philoxeroides* under different nutrient conditions, we studied the light response curve parameters of *A. philoxeroides*. Photosynthetic parameters, such as maximum photosynthetic rate (Pn_max_), light compensation point (LCP) and light saturation point (LSP), are important scientific basis for rapid growth of plants [62-64]. Light saturation point (LSP) can reflected the adaptability of plants to strong light; the lower the light compensation point (LCP) is, the better the normal photosynthesis is under the weak light and the maximum net photosynthetic rate (Pn_max_) reflects the utilization ability of *A. philoxeroides* to strong light under different treatments. In our study, among the three kinds of nutrient, the Pn_max_, LCP and LSP of the *A. philoxeroides* seedlings in medium planting density were the largest, indicating that under this treatment, *A. philoxeroides* had the strongest utilization ability and adaptability to glare, and under high planting density, *A. philoxeroides* has better capability to utilize weak light. Moreover, the photosynthetic parameters of the plant increased with the increasing nutrient levels. That is similar to previous studies [65, 66]. It shows that the increase of nutrient will enhance the utilization ability of *A. philoxeroides* to strong light. This suggests that at higher nutrient levels, higher light intensities are required to produce more biomass, may be that is why the *A. philoxeroides* has higher photosynthetic rate but the biomass is lower than the medium nutrient level. Under the same nutrient level, the increase of planting density resulted in the decrease of Pn_max_, LCP and LSP and there were significant differences among the three planting densities. The results suggested that the increase of planting density decreased the Pn_max_ of *A. philoxeroides* and its ability to use strong light and its adaptability. Under the low nutrient level, there was no obvious difference in the LCP between the medium and low planting density, which indicated that under the low nutrient level, the high planting density *A. philoxeroides* had more obvious photosynthetic ability at low light. But at the medium and high nutrient levels, the planting density had more obvious influence on the LCP of *A. philoxeroides*, and the effects became more obvious with the increase of nutrient level.

In conclusion, our study showed that at the three nutrient levels, the SLA, photosynthetic rate and total chlorophyll content of *A. philoxeroides* at medium planting density were the highest. What’s more, although the biomass of single plant, SLA, photosynthetic rate and the content of Chlorophyll reduced with the increase of planting density, the biomass of whole pot tended to increase. These attributes may increase the competitive dominance of *A. philoxeroides* and could help the *A. philoxeroides* population develop into a monodominant community.

